# Constructed Synchronized Multimodal Signalling in Great Bowerbirds

**DOI:** 10.1101/2022.10.20.513115

**Authors:** John A. Endler, Selina Meehan, Aida Rodrigues

## Abstract

Great bowerbird males build bowers for attracting females for mating. Bowers consist of a thatched twig tunnel (avenue) which opens onto two flat courts covered with objects. Male displays on a court are seen by a female from within the avenue. She sees and hears displays through the avenue entrance but can only see the male’s head and objects in his bill as it passes repeatedly across the entrance. We investigated bower acoustic properties by playing standard sounds from multiple court positions and recorded the resulting sounds at the female’s typical avenue head position within the avenue. Bower geometry significantly affects both his acoustic and visual display components and physically synchronizes them as he repeatedly moves in and out of the female’s view. Consequently, complex neural circuitry is unnecessary for linking sound to vision. Experimentally removing bower objects shows that objects significantly increase higher frequencies, hence bandwidth and loudness received inside the avenue. Great Bowerbird bowers produce a synchronized multimodal signal to females which may increase male attractiveness more than if a single sensory mode were used. This multimodal signal is unusual in that it is constructed rather than being part of the body and synchronized as a result of physics rather than neurons.

**Summary Statement:** Bowerbird bower geometry jointly effects both visual and auditory signal components and synchronizes them without the need for additional neural circuitry.

## Introduction

Multimodal signalling comprises components transmitted in more than one sensory mode and has been found in most animal groups [Mitoyen et al 2019]. Various functional, sensory, and behavioural functions, and consequent evolutionary advantages, have been attributed to multimodal signalling both within and among species [Hebets and Papaj 2005; Mitoyen et al 2019; Partan and Marler 2005; Ronald et al 2012]. Here we describe a case of multimodal sexual signalling which is affected by and synchronized by the geometry and composition of a construction by the male rather than being part of his body.

Male bowerbirds build a bower to attract females for mating [Frith and Frith 2004]. Great Bowerbirds (*Ptilonorhynchus* = *Chlamydera nuchalis*) build a bower consisting of a 60-100cm thatched stick tunnel (the avenue) opening onto two courts (Figure 1). Courts and avenues are constructed by the male and covered with stones, bones and bleached snail shells, and coloured objects (mostly green and red) are placed on the sides of the courts, close to the entrance [Endler et al 2014]. Objects are also placed in a depression in the middle of the avenue floor (Figure 1A). Male courtship takes place on a court near the entrance while the female watches from inside the avenue [Frith & Frith 2004]. She sees a complex visual display but can only see the male’s head, with or without coloured objects in his bill, whenever it passes within her field of view through the avenue entrance [Endler et al 2014].

**Figure 1.**
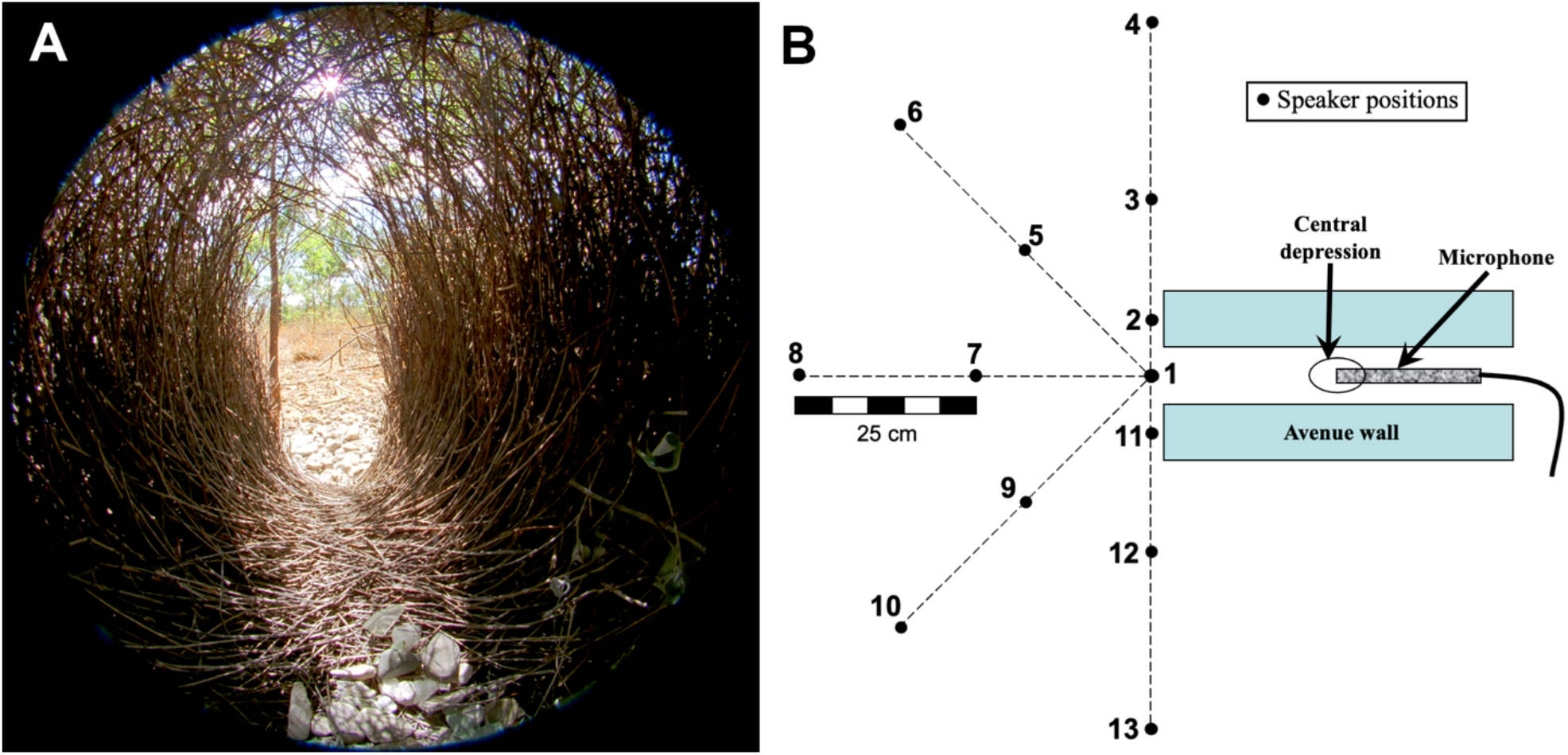
Great bowerbird typical bower structure. (**A**) Female great bowerbird’s view through the bower avenue, a small part of the bower court, and the area beyond the court, as shown in a hemispherical photograph with the “fisheye” lens held at the female’s eye level. Note the objects in the central depression within the avenue and on the court beyond the opening. She only sees the male’s head and a proffered object when his head passes across the avenue entrance. (**B**) Arrangement of the playback positions (1-13) on the main court and the position of the microphone within the avenue. At each position, the speaker was placed facing directly towards the avenue entrance (towards position 1). The photo (**A**) was taken where the tip of the microphone is shown in (**B**) facing towards positions 1,7,8.

Males produce a continuous vocalization during the display as they move their head in and out of the female’s field of view [Frith & Frith 2004, Endler et al 2014; Endler pers. obs, Figure 1]. We investigated how the bower structure and the dynamic position of the male during his visual display causes modulation and alteration of his vocalizations as received at the female’s head position within the avenue, because it can affect sound as well as light.

## Materials and Methods

Bowerbird males make a characteristic set of sounds (Figure 2A-F) at the bower court when actively displaying to a female in the bower avenue (Figure 1). If a female does not leave during the display, he walks around to the other avenue entrance, enters, and mates with the female within the avenue, continuing vocalizations until mating. The sounds do not guarantee a mating although longer bouts are more likely to be successful.

**Figure 2.**
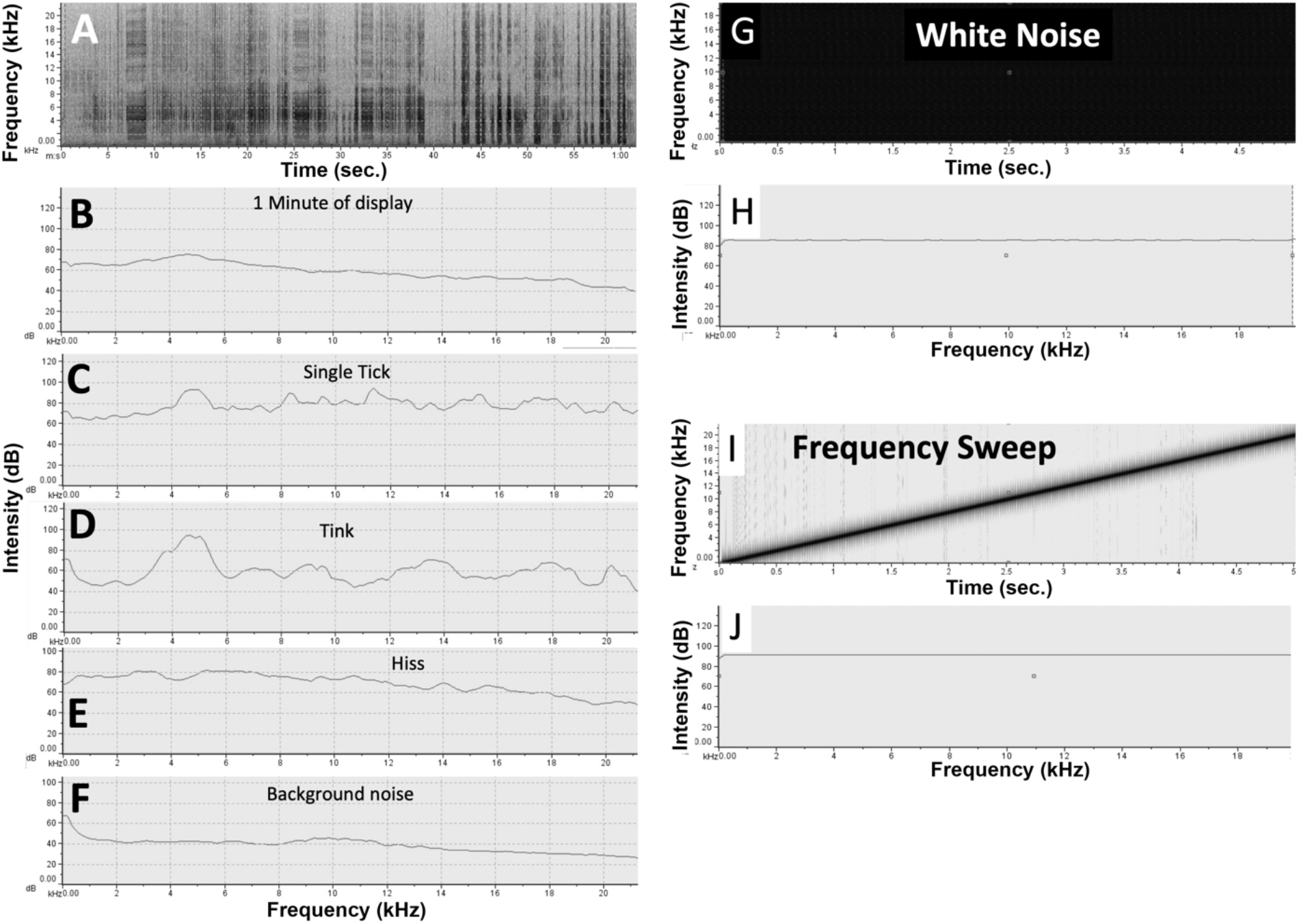
natural (**A-F**) and experimental (**G-J**) sounds. (**A**) spectrograph of characteristic sounds made by a great bowerbird male displaying on a court to a female inside the bower (scale 0-21 kHz by 0-60 sec.). (**B-F)** Sound spectra of sounds isolated from the recording (0-100 or 120 dB by 0-21 kHz). The background noise (**F**) was extracted from a quiet period in the same recording. Sounds made during displays are 20-40db above the background noise. (**G**) spectrograph white noise, a mixture of many frequencies (0 – 20 kHz by 5 seconds). (**H**) sound spectrum of (**G**) (0 – 120 db by 0 – 20 kHz). (**I**) Frequency sweep spectrograph. (**J**) sound spectrum of sweep (same scale as **H**). Playback spectra are flat, allowing simple estimates of sound properties of the bowers when recorded in the avenue.

In order to obtain fixed parameters for sound playbacks we recorded the display sounds at a bower on our Dreghorn Field Site, Queensland, Australia, with a Zoom recorder and microphone next to the bower at the same time as a motion-activated video camera recorded displays (as in Endler et al 2014). Figure 2A-F shows the sound properties of all sounds and each kind of sound extracted and analysed with RavenPro 1.5 software. There are two short components, ticks and tinks. Ticks are bursts of white noise repeated through much of the display whilst tinks are single almost bell like chirps and are much less frequent. The hiss is longer and shows as a rectangle in the spectrograph. These sounds have fairly flat spectra. Multiple recordings of display sounds by others (Xeno-Canto Foundation 2022) show the same spectral properties, and also include mimicry of other species. These are the basis of our standard sound parameters.

To provide a standard sound for playbacks at the Dreghorn bowers we constructed two different kinds of sounds using MATLAB 2021b with the same frequency range found in natural auditory displays, but flatter because we wanted to investigate the direct effects of the bower on the input sounds. One was a uniform mixture of frequencies (white noise) and the other was a frequency sweep (Figure 2G-J). The purpose was to play known replicable sounds and record what arrives at the female’s head position.

The sound files were played back from various positions on the main court (Figure 1B) using a Marantz PMD661 digital recorder resting end on the main court, with its speaker always facing towards position 1. The main court is the court where the male displayed most often [Endler et al 2014]. Playback positions 1 to 13 (Figure 1B) were designed to explore the effects of distance and angle to the avenue entrance. The playback volume was set just below the recording clipping level at position 1 and retained for all bowers.

Recordings of the played sounds were made with an AudioTechnica AT8035 microphone connected to a second Marantz PMD661 recorder. The microphone was placed in the avenue facing the main court with the tip centred over the avenue’s central depression, at the approximate position and height of the female’s head (Figure 1) in video recordings [Endler et al 2014]; about 4cm below the avenue roof. Recordings were also made in open areas away from a bower or vegetation with the same geometry to estimate the effects of the playback, speaker and microphone properties on the received spectra. Recordings were converted to frequency spectra (dB vs sound frequency), using the MATLAB 2021b SoundSpectrum function, and used in subsequent analysis. The artificial sound spectra were flatter than the bowerbird sounds (Figure 2) in order to explore the auditory properties of the bower avenue at each frequency rather than being a mimic of natural sounds, and also to make up for the relatively weaker high frequency response of speaker and microphone. Spectra were analysed by GAM (General Additive Models) and results plotted using the R (R Core Team 2022) packages mgcv (function gam, Wood 2022) and itsadug (plot_smooth, van Rij et al 2022).

We performed two experiments at natural bowers to obtain data on what a female bowerbird could hear while she stands inside the bower avenue looking at and listening to the male display. The ornament experiment examined the acoustic effects of removing objects from the bower that were previously placed on the bower by the male bowerbird. The directionality experiment explored the acoustic directional properties of the avenue on undisturbed bowers. Both experiments were performed on active bowers, indicated by the presence of green fruits beside the avenue entrance, some red objects off the courts and a neatly constructed avenue. We are making no assumptions about the utility of the sounds nor are we assuming that the sound properties are intentional, bower structure could easily affect bowerbird sounds; our intention is to investigate how much effect the bower structure has on sound received by the female.

Ornament Experiment. In November 2015 we recorded playbacks of the white noise sound (Figure 2G,H) from speaker positions 1, 7 and 8 (Figure 1B) at four Dreghorn bowers D02, D15, D32, D43 which had abundant bleached snail shells, and open areas. Fraction shells varies greatly among bowers and locations; we chose these bowers to minimize shell cover variation. Each bower was subject to 5 treatments, each followed immediately by recording with the speaker at positions 7 and 8. The experimental treatments were as follows: Nat: Natural or undisturbed bower. CtO: Objects in court only, removed snail shells and bones (when present) from the bower avenue. CDO: Objects in central avenue depression only, replaced objects in the avenue, removed objects from the courts. Most avenue objects are placed by the male in the depression in the centre of the avenue. FSS: filled snail shells, filled court shells with sand and dough (flour and water) to change their sound properties and then put them back on the court. WSS: without snail shells or other objects (e.g. bones, stones), all objects removed. We did not disturb the bower except for the object manipulations and then carefully placed the microphone in the avenue and the speaker in the three court positions (sequentially) for playback. After all treatments (about 1.5hr) the removed objects (including shells emptied of sand and dough) were placed near the bower. The bowerbird replaced them on the bower within a day or two, as in previous object manipulation experiments (Endler et al 2010, Kelley & Endler 2012). We analysed the effects of the ornaments with the speaker placed at 50cm, 25cm and 0 cm from the entrance. Even at 0 cm some ornaments extend into the avenue. The male places his head in and out of the female’s field of view from 10 to about 25 cm from the avenue entrance.

Directionality Experiment. In August 2021 we recorded playbacks of frequency sweeps (Figure 2 I,J) from all 13 positions (Figure 1B) at five active bowers D01, D02, D34, D44, D45 and three open areas to control for speaker and microphone properties. We did not disturb the bower except for carefully placing the microphone in the avenue (tripod at the back entrance) and the speaker at each position. We used a frequency sweep rather than white noise because we wanted closer control over amplitude than is possible with a mixture of frequencies with the same frequency range (2Hz - 20kHz). In order to allow for minor differences in bower size and equipment positioning we subtracted the data from position 1 (Figure 1A) from the data from the other positions in the same bower or open area. This yields the relative effects on sound arriving at the avenue entrance from various distances and angles. This is a measure of sound attenuation geometry.

## Results

Figure 3 shows the results of a GAM analysis on the effects of treatment on the frequency spectra for 25cm between the speaker and the avenue entrance. The results of the GAM analysis were similar for all 3 distances from position 1 (Figure S1). Effects of frequency (spectral shape) and interaction between frequency and treatment were both highly significant (P<10^−16^) for each distance. The r^2^ were 0.949, 0.945, and 0.949 and the deviance explained were 95, 94.6, and 94.9% for 50, 25 and 0cm from the avenue entrance, respectively (see online supplement tables S2 and S3 for analyses).

**Figure 3.**
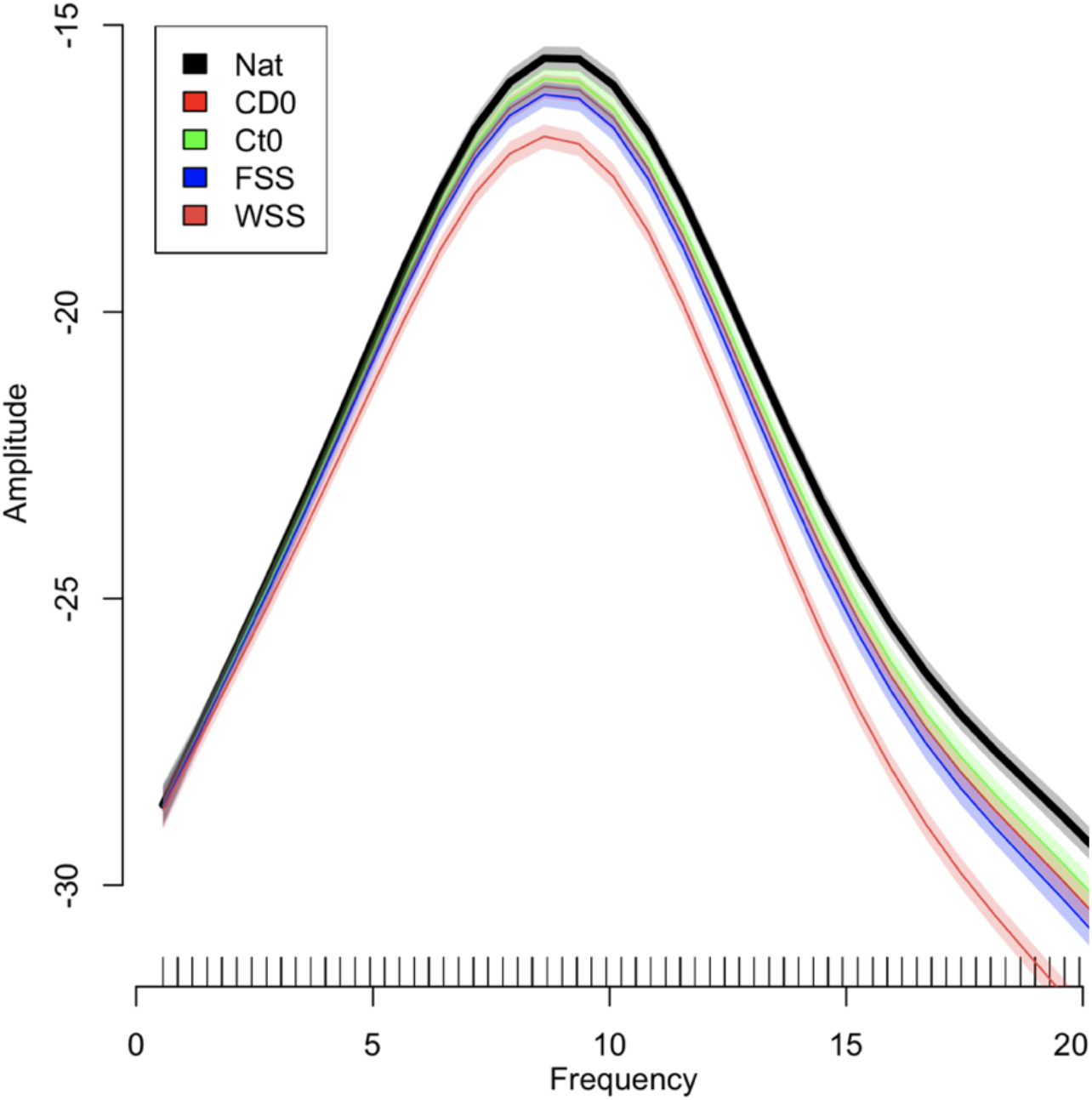
GAM analysis of sound spectra in each treatment with playback at 25cm from main court avenue entrance. Frequency is kHz and amplitude is in dB units. Lines are the GAM estimates and the shaded areas around them are their estimated errors. Differences are significant when the error zones do not overlap. The black line and grey shading indicate the undisturbed bowers and the others are for various ornaments removed or modified. Removing all objects (WSS) has the largest effect, but other removals (CDO, CtO, FSS) do not differ significantly from each another.

For all three distances there is a clear effect of both amplitude and spectral shape from ornaments, as shown by the highly significant shape x treatment interaction effect. The ornaments increase sound amplitudes significantly at above around 5kHz and are particularly effective in increasing amplitudes above around 10kHz. The material on the court and avenue are very important in maintaining relatively more higher frequencies as shown by the effects of removing them.

Removing either the court or avenue shells separately or filling the shells (CDO, CtO, FSS) has a smaller effect than removing everything (WSS). Unexpectedly, there was little difference between the effects of CDO, CtO and FSS (FIgure 3). This suggests that the ornaments inside the avenue as well as the court are important in maintaining the broadband sound quality for the female inside the avenue. This is expected if the effects of ornaments are due to efficient reflection of higher sound frequencies both outside and inside the avenue.

Figures 4 and S2 show the total sound intensity of the same sound originating from various angles and distances from the avenue entrance for bowers (Figure 4B) and for the same geometry of speakers and microphone in the absence of bowers (Figure 4A). It is clear that the avenue has a very large effect on the geometry of sound intensity. It is highest if the sound comes from the avenue’s long axis and declines very rapidly when the sound comes from increasing angles from the avenue long axis. The effect is greatest between 0 and 10cm from the entrance which is where the male bowerbird displays to the female (4-12db reduction). Although the microphone has a directional response (Figure 4A) the sound intensity geometry is profoundly affected by the avenue, becoming spatially restricted to a narrow zone aligned with the avenue axis (Figure 4B). Even small side-to-side movements of his head in and out of the avenue axis can yield 5-6 dB intensity variation.

**Figure 4.**
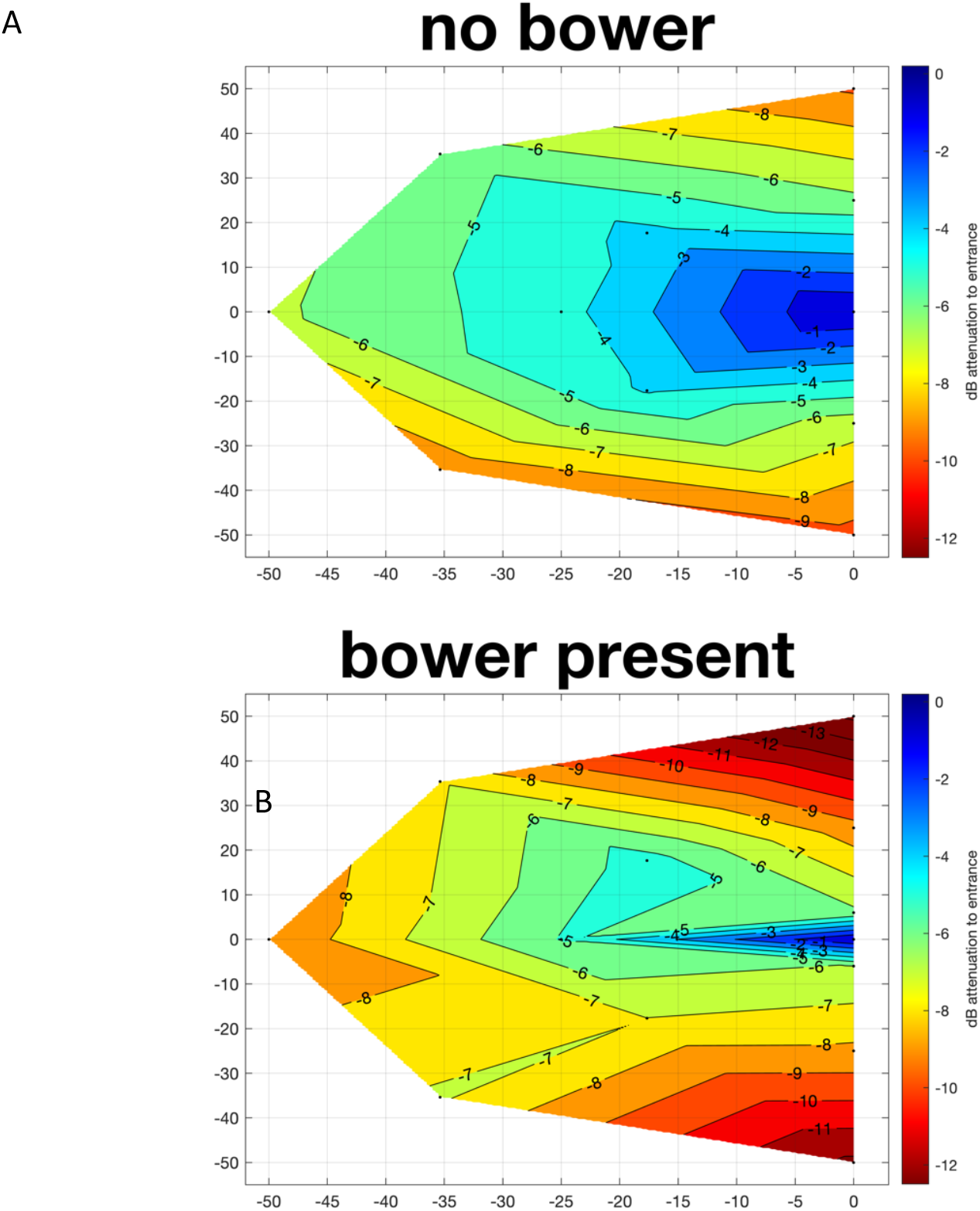
Sound intensity (dB) relative to position 1 (Figure 1B) as a function of the playback speaker at various distances and angles from position 1. A: Data from three open areas. B: Data from five bowers. The graph origin (0,0) (position 1) corresponds to the avenue entrance in B. Distances on the main court in cm, negative to the left of position 1. Contour labels in dB; note identical scales in A and B. More red means sound is more attenuated. Irregularities result from minor variation in position and speaker angle and objects on the court. A shows the inherent directionality of the microphone but it is clearly not as directional as the bower effects (B).

## Discussion

Bower avenue geometry and the ornaments placed on the bower clearly effect the composition of the auditory component of the Great Bowerbird display. The ornaments enhance the higher frequencies of sounds reaching the avenue centre, making the input spectrum broader (flatter) than it would be without the reflections from the ornaments (Figure 3). This better matches the auditory display spectrum (Figure 2) presumably increasing intensity and auditory contrast. The avenue shape means that sounds originating outside the avenue only reach the female in the avenue centre at high amplitude when the source is directly in line with the avenue long axis (Figure 4) as happens intermittently as the male’s head and/or proffered objects move in and out of the avenue axis, just as in the visual part of the display. Geometry ensures synchronization of sound with vision.

Although the sound intensity enhancement of the ornaments is significant we do not know what higher frequencies bowerbirds can hear. Passerines can generally hear up to about 10kHz and sensitivity declines rapidly above that even if they emit higher frequency sounds (Köppl 2022). Gleich (Gleich et al 2005; Gleich and Langemann 2011) showed that there is a good relationship between body size, basilar papilla length and the optimum and maximum audible sound frequency for all birds (with Strigiforms being much better than the other groups) and provided equations. These equations seriously underestimate the maximum for hummingbirds yielding 8.5kHz instead of the observed 15kHz for *Oreotrochilus chomborazo* (Duque et al 2020). The equations for a bird with a Great Bowerbird mass (200gm) is predicted to be about 6kHz, but given the very broad bandwidth of their tick, this may also be an underestimate. We explored the possible effects of a lower maximum by repeating the GAM analysis of the spectra after cutting it off above 10.5 and 5.5kHz, see Table 1. The relevant test is for the sound frequency by treatment interaction, where treatments are the five manipulations of the ornaments. It is clear that the ornaments have a significant effect at all distances when the highest frequency is cutoff at 10.5kHz. At a 5.5kHz cutoff It is still significant, but smaller at 0 and 30 cm from the entrance but not at 50cm. Consequently the effects of ornaments increasing bandwidth could be audible, and at the very least increasing the total intensity variation during the display.

**Table 1.** results of GAM on the received sound frequency from source at 3 distances and either the entire bandwidth (to 20 kHz) or truncated to 10.5 and 5.5kHz.

| Distance | to 20kHz | to 10.5kHz | to 5.5 |
| --- | --- | --- | --- |
| kHz |  |  |  |
| to Avenue | P, Frequency x Treatment |  |  |
| 50cm | 2.00E-16 | 2.00E-16 | 0.088 |
| 25cm | 2.00E-16 | 2.00E-16 | 0.017 |
| 0cm | 2.00E-16 | 1.89E-05 | 1.58E-05 |
|  | P, |  |  |
|  | Frequency |  |  |
| 50cm | 2.00E-16 | 2.00E-16 | 2.00E-16 |
| 25cm | 2.00E-16 | 2.00E-16 | 2.00E-16 |
| 0cm | 2.00E-16 | 2.00E-16 | 2.00E-16 |
|  | Deviance Explained |  |  |
| 50cm | 95% | 96.2% | 92.9% |
| 25cm | 94.6% | 95.8% | 90.6% |
| 0cm | 94.9% | 96.2% | 92.2% |
|  | N |  |  |
| each | 1311 | 646 | 342 |

Great bowerbird bowers vary with location in the number of bleached snail shells they use, even over a distance of a few km; this study was performed in an area where shells were most often used on bowers in order to minimise inter-bower variation of ornaments. If increasing bandwidth is important, bowers where few shells are available could use other possible ways of increasing sound bandwidth such as using more flat stones, which are ubiquitous in great bowerbird bowers and have a fairly flat reflection spectrum to at least 15 kHz (Isele et al 2009). Spotted Bowerbird (*P. maculatus*) bowers also vary in the numbers of shells used from place to place (Madden 2006) and where used they often make up almost all of the court, unlike Great bowerbird courts where snails rarely cover all the court. Great bowerbird avenues are made of densely thatched twigs and sticks whereas Spotted bowerbird avenues are made of robust grass stems and are shorter and wider than Great bowerbird avenues (Frith & Frith 2004). As a consequence do more snails on the spotted bowerbird courts improve the sound bandwidth enough to make up for a softer and wider bower (less sound focusing and less sound reflection)? Also, are avenues of both species more dense and longer in areas where there are fewer snails available or weaker snail choice? Other bowerbird species do not or use few snails or other hard objects in their bowers (Frith & Frith 2004; Endler et al 2005); how does that effect their auditory sound components?

Male bowerbird displays include a continuous set of sounds (Figure 2A) whilst moving his head back and forth across the avenue entrance and long axis. The physics of sound interacting with the bower means that the amplitude received by the female should show large excursions with the male’s head movements (Figure 4B, large intensity differences at less than 10cm from the avenue entrance). Consequently, the auditory and visual components are synchronised, alternatively becoming salient and minimal. Unlike synchronised multimodal displays in other species, which depend entirely on synchronized motor behaviour and specialised neural circuitry, here all the male bowerbird has to do is to move his head across the avenue entrance as part of his visual display, and ensure that the decorations provide the visual stimuli and enhance the auditory stimuli bandwidth.

We do not know if the sounds are important in mating, but if not why would they make them? The hiss component is also used when other males visit the bower so it may have a dual purpose. The sound would be continuous to other males, who would not be in the avenue, but fluctuating in amplitude to a female in the avenue. It might be hard to demonstrate the value of the sound component of the male bowerbird display because the physics of sound interacting with the bower structure and shape, in combination with the male’s head movement across the avenue entrance, cause the amplitude fluctuations, and more objects are more attractive, possibly for visual but possibly for both visual and auditory reasons.

Male bowerbirds constantly move in and out of their bower avenue, going inside, looking out, then moving objects on the court, and repeating this for many days. This behaviour serves to maintain and improve the visual pattern (Endler et al 2014, Kelley and Endler 2012). Given that rival males visit the owner’s bower periodically and both birds hiss, it i conceivable that the owner male could modify the sound properties after listening to the rival’s hiss when he is inside the avenue testing the visual effects. This needs testing.

Constructed devices to intensify and modify sound signals are also known in mole crickets (Daws et al 1996), tree crickets (Mhatre et al 2017), and frogs (Muñoz and Halfwerk 2022), but these are not multimodal. Unlike the bowerbirds these are all examples of constructions augmenting signal emission and transmission from the male whereas in bowerbirds the construction affects the receiver (females). Hummingbirds also use physics to synchronize visual and vocal signals, taking advantage of the significantly directional gorget colours which flash during the motion and auditory display (Duque et al 2022; Hogan and Stoddard 2018), in this case the gorget flash is passive and the sounds and flight patterns are active. These display movements are likely to be energetically very costly. However, having a physics-based flash saves extra energy which might otherwise be needed for actively moving plumage. What is particularly unusual about the great bowerbird multimodal display is that it is actively constructed by the male in order to affect female reception of the signal, and the consequence of the constructed bower is a synchronized visual and auditory display. No extra energy or neural circuitry is required for the auditory synchronization. All of these examples show the value of accounting for the geometry of mate signalling (Echeverri et al 2021).

If synchronization is expensive, how often are multimodal signals in other species unsynchronised or not tightly synchronized owing to the costs of motor and muscular synchronization? Are species living in energy-poor habitats less likely to use synchronized multimodal signals than those living in energy-rich habitats?

## Acknowledgements and Funding

We thank Jack Bradbury, Robert Dooling, Laura Kelley, Joah Madden and Steve Rothstein for very helpful suggestions and comments on the manuscript. This research was supported by fees for being the Editor-in-Chief of Evolutionary Ecology, Deakin University CIE funding, Holsworth Foundation, and JAE’s personal funds.

